# Designing an occupancy framework to monitor an endemic rainforest duck: Methodological and modelling considerations

**DOI:** 10.1101/2020.02.03.931543

**Authors:** Shah Nawaz Jelil, Murchana Parasar, Laura Cancino, Kimberly Cook

## Abstract

Understanding species trend, decline or growth, is vital to further conservation efforts. Species-habitat relationship studies are equally important for conservation as it helps in understanding the habitat a particular species depends upon, i.e. habitat conservation. However, rare and endemic species are inherently difficult to study and occupancy models are especially useful in such cases. We conducted the first detection, non-detection survey for the white winged duck in Dehing Patkai Wildlife Sanctuary, India to assess site occupancy and test habitat factors that explain its occupancy. We found that white winged duck occupancy was low (0.27 ± 0.21 SE) and detection probability was 0.44 ± 0.30 SE. We found that increasing tree richness and decreasing elevation increased species occupancy. Detection probability was influenced by our effort in that detection increased with increasing number of survey hours. Using two standard approaches, we estimated the optimal number of sites and replicate surveys for future occupancy studies. We further present considerations for future surveys. Considering the sporadic and fragmented information available, we recommend long-term ecological research to better understand the present and future population trends of the species.

## Introduction

Estimating occurrence and abundance of focal species remains a fundamental objective in initiating or furthering conservation efforts. This becomes particularly vital when dealing with species which are endemic, elusive and/or have restricted ranges. The white winged duck *Asarcornis scutulata* is one such species. It is one of the largest duck species, which are dependent on small wetlands amidst lowland areas of tropical moist forests in South-East Asia (Green, 1993). The species was previously widely recorded from northeast India, Bangladesh and Myanmar, Thailand, Laos, Vietnam, Cambodia, peninsular Malaysia and Indonesia (Green, 1993). However, recent studies estimate just over 400 individuals to be present in India (Choudhury, 2002) and this represents more than a third of the global population (Sharma et al. 2015). In India, the species is distributed in the lowland tropical rainforests of Assam and Arunachal Pradesh (Talukdar, 1992; Choudhury 2000; Rahmani and Islam 2008) and in Meghalaya (Hume 1890; Choudhury, 2002). The primary factor for the white winged duck decline has been assigned to the disturbance and destruction of the primary rainforest habitat all across its distribution range (Mackenzie and Kear, 1976). Decrease in total area of suitable habitat, isolation of different groups of ducks, increase in disturbance within forests, increased risk from illegal hunting and risk of pollution of available waters by industrialized small towns, wood mills and tea estate waste have been attributed as the major reasons of the species decline in Assam (Mackenzie and Kear, 1976). A major natural event which affected the duck and the primary rainforests in the state of Assam was the 8.5 magnitude Assam-Tibetan earthquake of 1950. This resulted in the inundation of important forest tracts of Sadiya to Lakhimpur North and the Dibrugarh Reserve Forest. These forests were swamped and largely destroyed. However, this was an exception in the already ongoing and incessant habitat transformation due to human activities. Subsequent cane cutting, hunting, buffalo grazing, fishing and flight netting only hastened the species decline (Mackenzie and Kear, 1976). Hence, it is not surprising that recent sightings have been scarce and largely opportunistic (Selvan et al. 2013; Sharma et al. 2015). The lack of information on its present status and dearth of ecological knowledge about the species and the habitats that they depend on endangers the species greatly. Not only is the current status of the bird unknown, but methods and analytical framework to assess current trends and monitor the species remain unclear and without agreement among scientists and field researchers. With this backdrop, we explored the use of occupancy models to assess and monitor the species in the lowland remnant rainforests of Dehing Patkai Wildlife Sanctuary in Upper Assam, India. Occupancy models provide rigorous framework and hence have been widely used across species and habitats (Edwards et al. 2018). It was fitting to use an occupancy framework for our study as it requires lower number of survey sites than abundance surveys and hence are less expensive and labour intensive (Mackenzie et al. 2006; Edwards et al. 2018).

## Materials and methods

The Dehing Patkai Wildlife Sanctuary (Fig. 1), with an area of 119.9 km^2^, stretches in the Tinsukia and Dibrugarh districts of the state of Assam, India. Dehing Patkai consists of three main parts, viz. Soraipung (part of Upper Dehing), Jeypore Reserve Forest and Dirok Reserve Forest. The wildlife sanctuary is also a part of the Dehing Patkai Elephant Reserve. The forest is a part of an important bird area (IBA) IBA-the Upper Dehing West Complex. The habitat is tropical rainforest and described as Assam Valley tropical wet evergreen forest (Champion and Seth, 1968). The forest is characterized by a top canopy dominated by *Dipeterocarpus macrocarpus* reaching heights of 50 m, a middle canopy dominated by *Mesua ferrea* and *Vatica lancefolia* and undergrowth consisting of various shrubs such as *Saprosma ternatum, Livistonia jenkinsiana* and *Calamus erectus* (Kakati, 2004; Saikia and Devi, 2011).

**Fig. 1:**
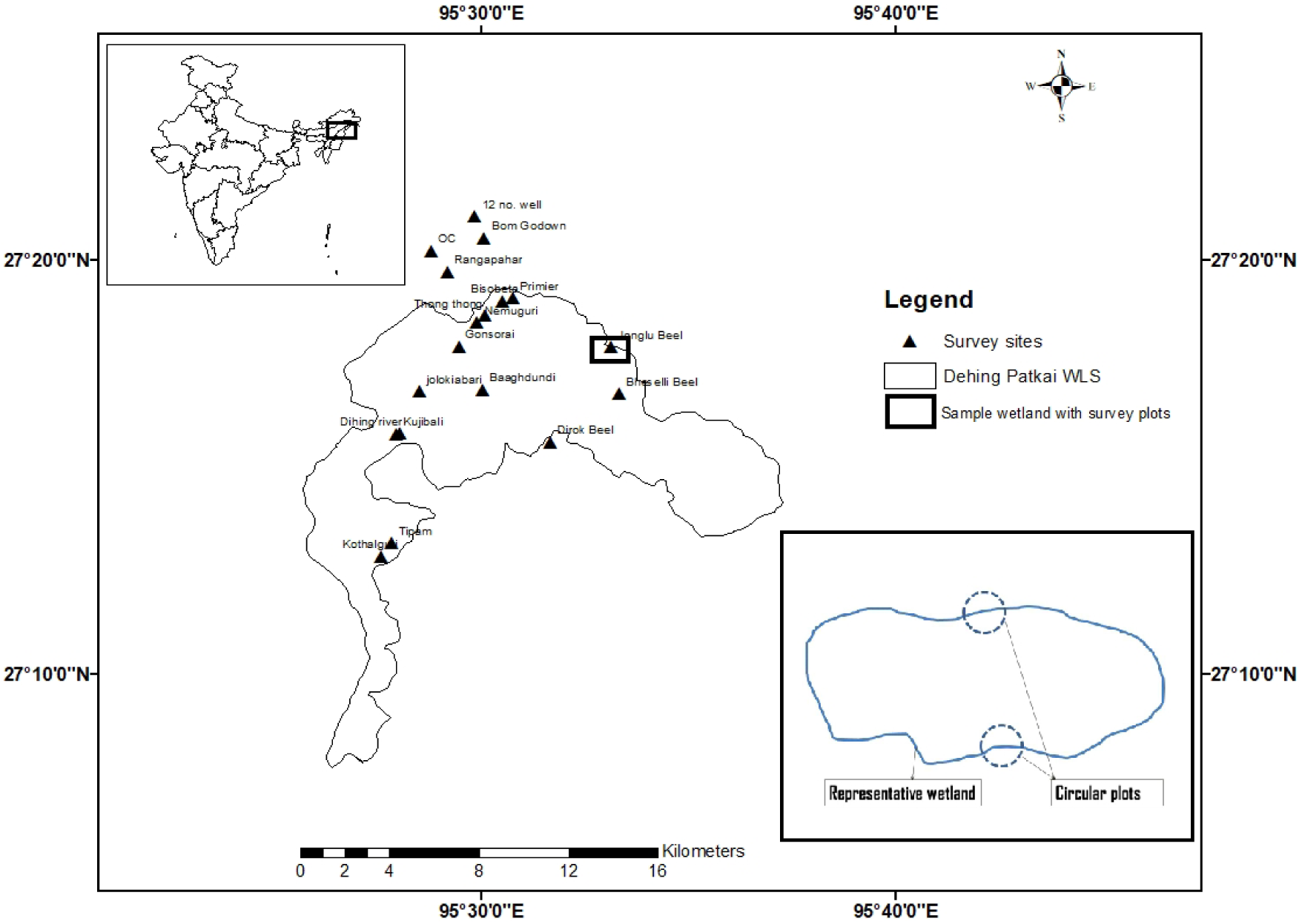
Map of Dehing Patkai Wildlife Sanctuary with surveyed wetlands.

Owing to the dearth of ecological knowledge of the species in the region, we initially conducted a recce survey from April to June, 2017 wherein we coordinated with the local forest department officials and local villagers to get acquainted with the wetlands with earlier records of the duck. The intention of the recce, especially the survey with the forest officials was to assess if the bird still used these wetlands. This however did not bias the selection of our wetlands for detection-non detection surveys. We surveyed 18 wetlands as these sites are naturally occurring discrete ponds and hence act as independent units (Mackenzie et al. 2003). We followed the standard design for site occupancy estimation where *s* number of sites is surveyed *K* times in one season; in our case it was the post-monsoon season of 2017. Surveys were carried out in wetlands located inside and within *c*. 10 km buffer from the wildlife sanctuary boundary. Our surveys were carried out twice within the pre-monsoon season and hence we had two replicates for each survey site. The initial survey was conducted in November 2017 and the second survey in December 2017. Each survey was carried out by three experienced observers in all the wetlands for a minimum of two hours (Table 1). Effort was measured as the number of hours spent in each of the replicate.

**Table 1:** Occupancy and detection covariates used in the occupancy models.

| Covariate type | Covariate | Type of variable | Code and code name used | Mean values and range |
| --- | --- | --- | --- | --- |
| Occupancy | Canopy cover (%) | Continuous | canopy | 73.22 (50–85) |
|  | Tree richness | Continuous | tree | 10.33 (5–16) |
|  | Elevation | Continuous | ele | 8872.38 (806–943) |
|  | Number of logs in wetland | Continuous | log | 8.05 (6–13) |
| Detection | Effort | Continuous | effort | 3.61 (2–6) |
|  | Survey occasions | Categorical | survey<br>P[1]: 10<br>P[2]: 01 | — |

## Site covariates

We established 10 m radius plots (314 m^2^) in all the surveyed wetlands to assess the number of trees, dominant tree species, percent canopy cover and presence of logs in streams. Considering the structure of the wetlands, two plots were established on either side. The plots were established in a manner so that half of the circle was in the wetland and half outside (Fig. 1). We then estimated the mean of all the parameters from the two plots in each of the wetland. We report mean canopy cover (%), tree richness, fallen log richness and elevation. These covariates were selected *a priori* because of their likely importance in driving white winged duck occupancy in the study area. Large mature trees such as Hollong *Dipterocarpus macrocarpus*, Teak *Tectona grandis* or Mekai *Shorea assamica* are seen as suitable habitat for the white winged duck (Mackenzie and Kear, 1976).

## Data analyses

We conducted single season occupancy models (MacKenzie et al. 2002; 2006) using PRESENCE (v. 12.6) (Hines 2006). We created the detection history matrix using two replicates and ran different combinations of covariate models for the species. The most parsimonious models were selected on the basis of Akaike Information Criterion (AIC) (Burnham and Anderson 2002) and estimates for occupancy (*ψ*) and detection probability (*p*) were obtained from the null model. The habitat data collected were input as occupancy covariates in models (Table 1). We modelled detection as a function of survey occasions and our effort (hours of survey). We later constructed linear models to check the effect of covariates on the species occupancy (*ψ*) using R (v. 3.4.3) (R Core Team 2017).

## Limitations and future surveys

Due to logistical constraints, we could conduct only limited number of surveys and hence using the occupancy modelling framework was the best fit. However, we acknowledge that minimising number of surveys may lead to incorrect estimation of *ψ* value with increased error. We explored two options/approaches to overcome this limitation. First, we estimated the optimal number of replicate surveys (*K*) needed and the number of survey sites (*s*) to achieve an occupancy estimate with a precision of 0.05. This was done following the framework provided by Mackenzie and Royle (2005) and also successfully applied by Edwards et al. (2018). We input our *ψ* and *p* estimates into Table 1 of Mackenzie and Royle (2005) to estimate *K* and used equation 3 to estimate *s* which is:

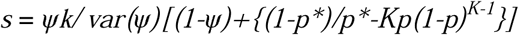

where, *s* = number of sites

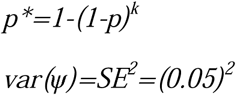

The other approach we used was recently proposed by Specht et al. (2017) which is specifically designed for species occurring at low abundances (*ψ* ≤ 0.3). For this, we input our *ψ* and *p* estimates into Table 2 of Specht et al. (2017) to estimate the optimum number of *K* (number of visits to survey sites) considering that the cost of repeated surveys remain the same.

## Results

We encountered two groups of white-winged ducks of cluster size 5 and 3 respectively in Bomgodown and Rangapahar wetlands. In OC and 12 no. well wetlands, we encountered individual ducks once during the second survey/replicate. We found that all wetlands were surrounded by large trees, *Shorea assamica, Vatica lancefolia, Dillenia indica, Mesua ferrea* and *Dipterocarpus macrocarpus*.

Occupancy and detection probabilities and their predictors: The proportion of site occupied by the white winged duck was 0.27 (± 0.21 SE; naïve occupancy = 0.16). Detection probability of the duck was 0.44 (± 0.30 SE). Although number of trees and number of fallen logs had a combined effect (Table 2), we found that tree richness positively affected and explained much of its occupancy (R^2^ = 0.89) (Table 3; Fig. 2). Number of fallen log affected negatively however this was negligible (R^2^ = −0.062) (Table 3; Fig. 3). Another factor affecting species occupancy was elevation. We found as elevation increased, white winged duck occupancy decreased (R^2^ = 0.41; Fig. 4). We found that that the probability of detection for the species was affected by our effort (R^2^ = 0.41; Fig. 5) and that the majority of the improvement over the null model was driven by *p*(effort). Detection probabilities increased with increasing effort. Hence, we suggest input of more effort to produce a more precise estimate with a substantially low standard error.

**Table 2:**
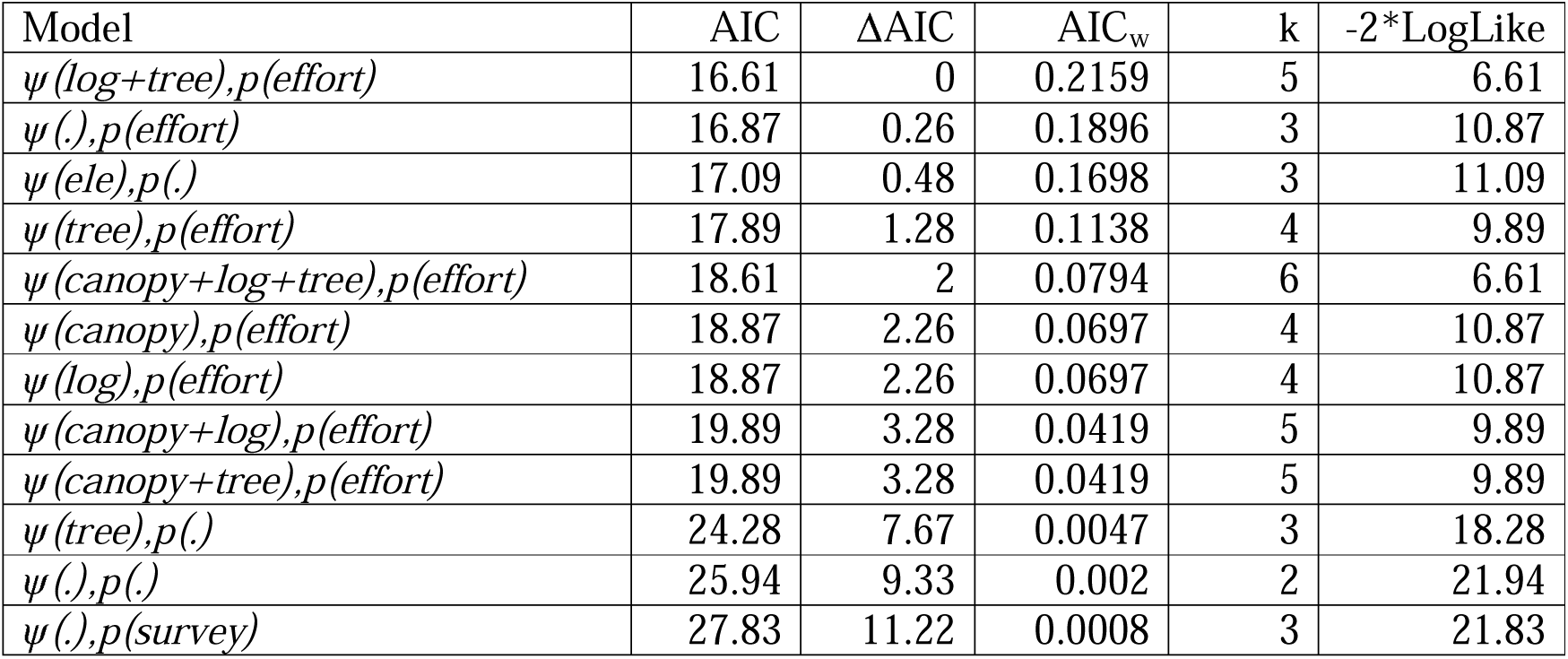
Table showing various combinations of occupancy models.

**Table 3:** Table showing the significance of the top covariates in explaining white winged duck occupancy (*ψ*) and detection (*p*)

| Variable | Coefficients | Estimate | SE | Multiple R<sup>2</sup> | Adjusted R<sup>2</sup> | p |
| --- | --- | --- | --- | --- | --- | --- |
| $\Psi$ | Number of trees | 0.09 | 0.007 | 0.90 | 0.89 | 2.807e-09 |
|  | Number of logs | -2.417e-05 | 7.673e-04 | 6.198e-05 | -0.062 | 0.97 |
|  | Elevation | -0.006 | 0.001 | 0.44 | 0.41 | 0.002 |
| $p$ | Effort | 0.14 | 0.04 | 0.44 | 0.41 | 0.002 |

**Fig. 2:**
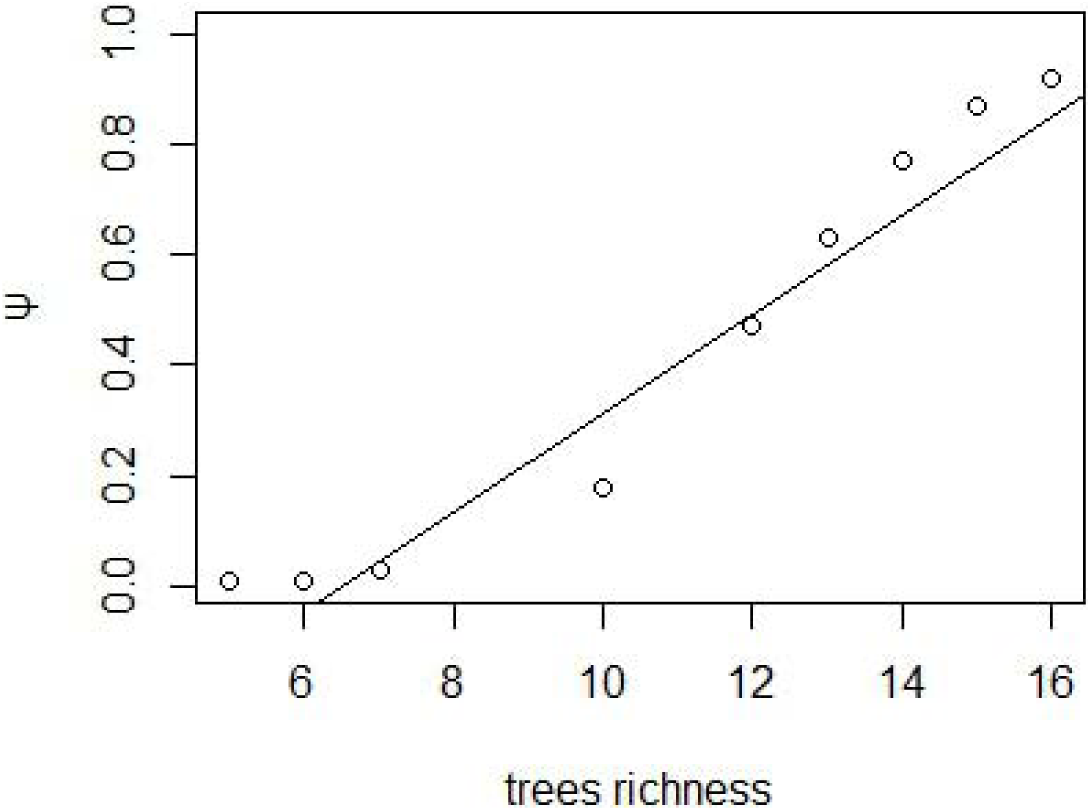
Effect of tree richness on occupancy of white winged duck.

**Fig. 3:**
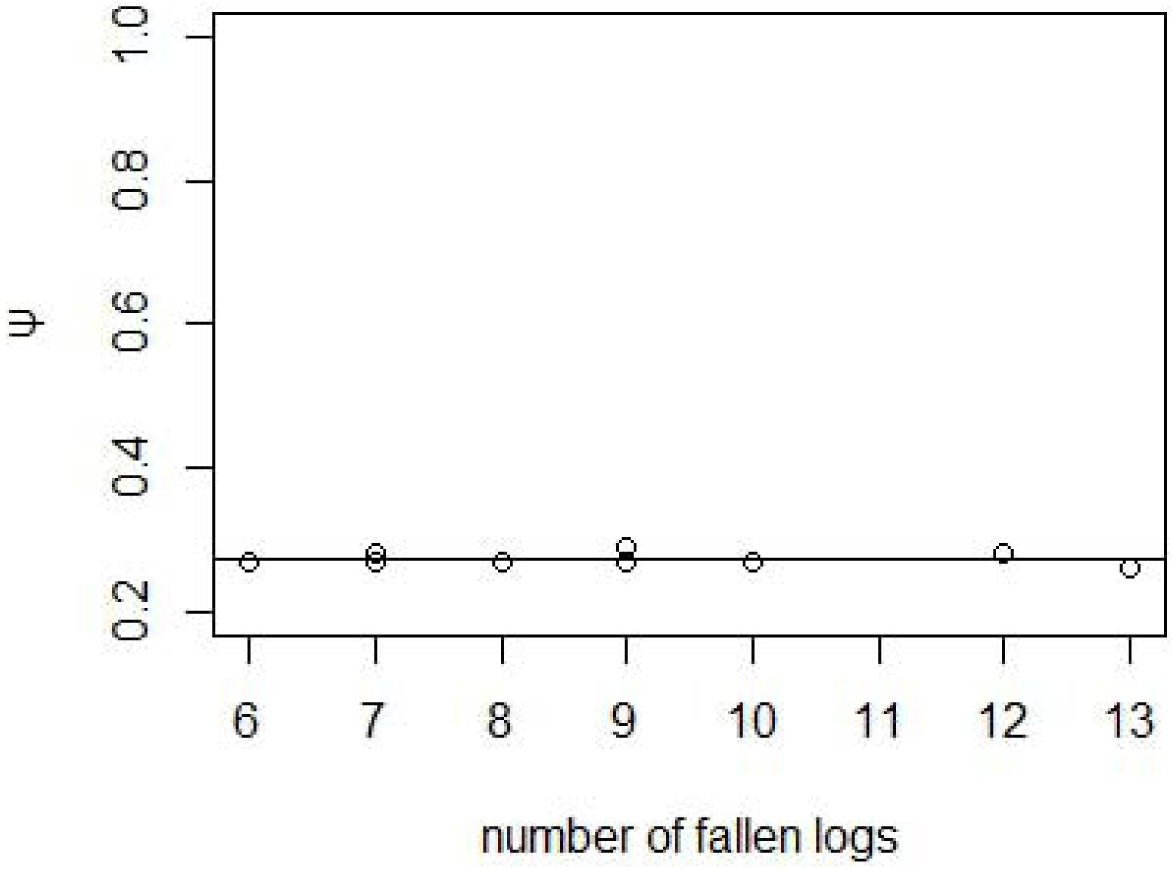
Effect of number of fallen logs on occupancy of white winged duck.

**Fig. 4:**
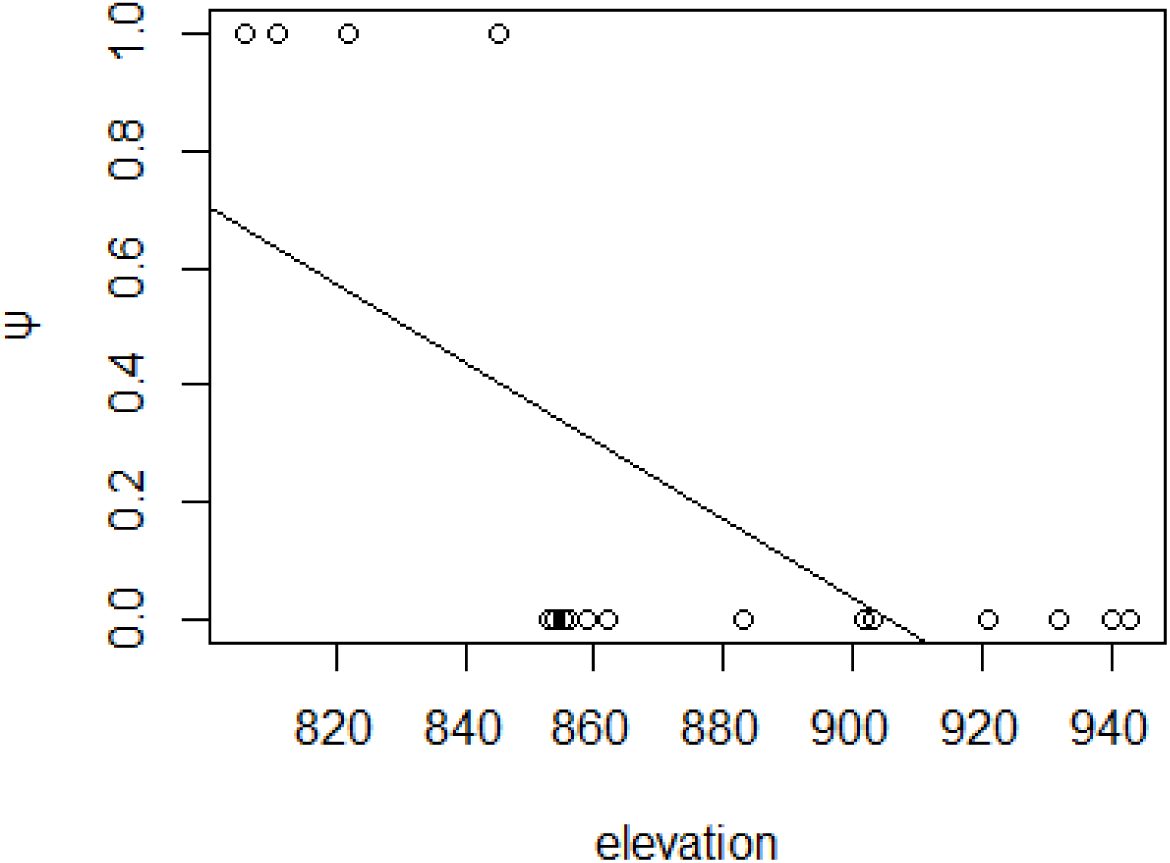
Effect of elevation on occupancy of white winged duck.

**Fig. 5:**
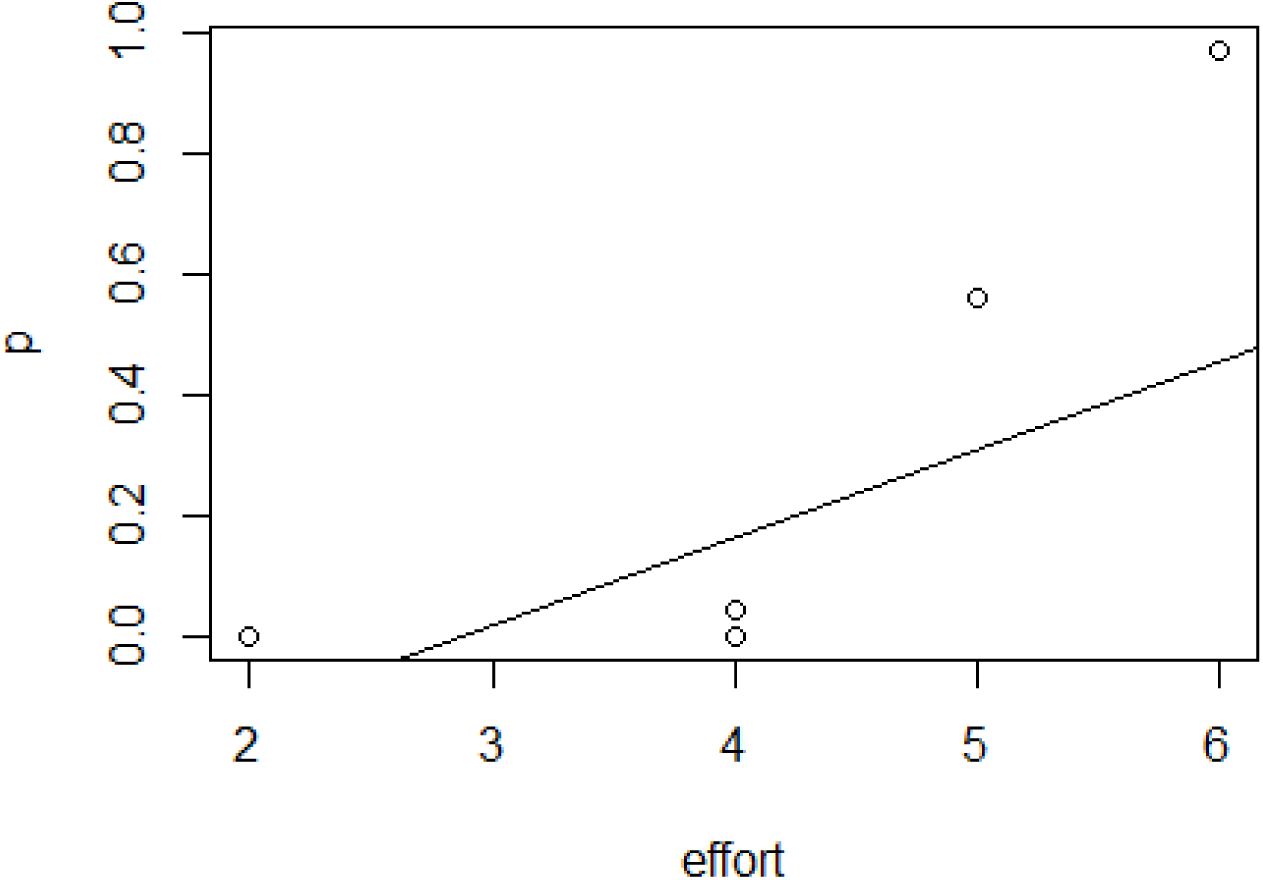
Effect of our field survey effort on detection probability of white winged duck.

**Fig. 6:**
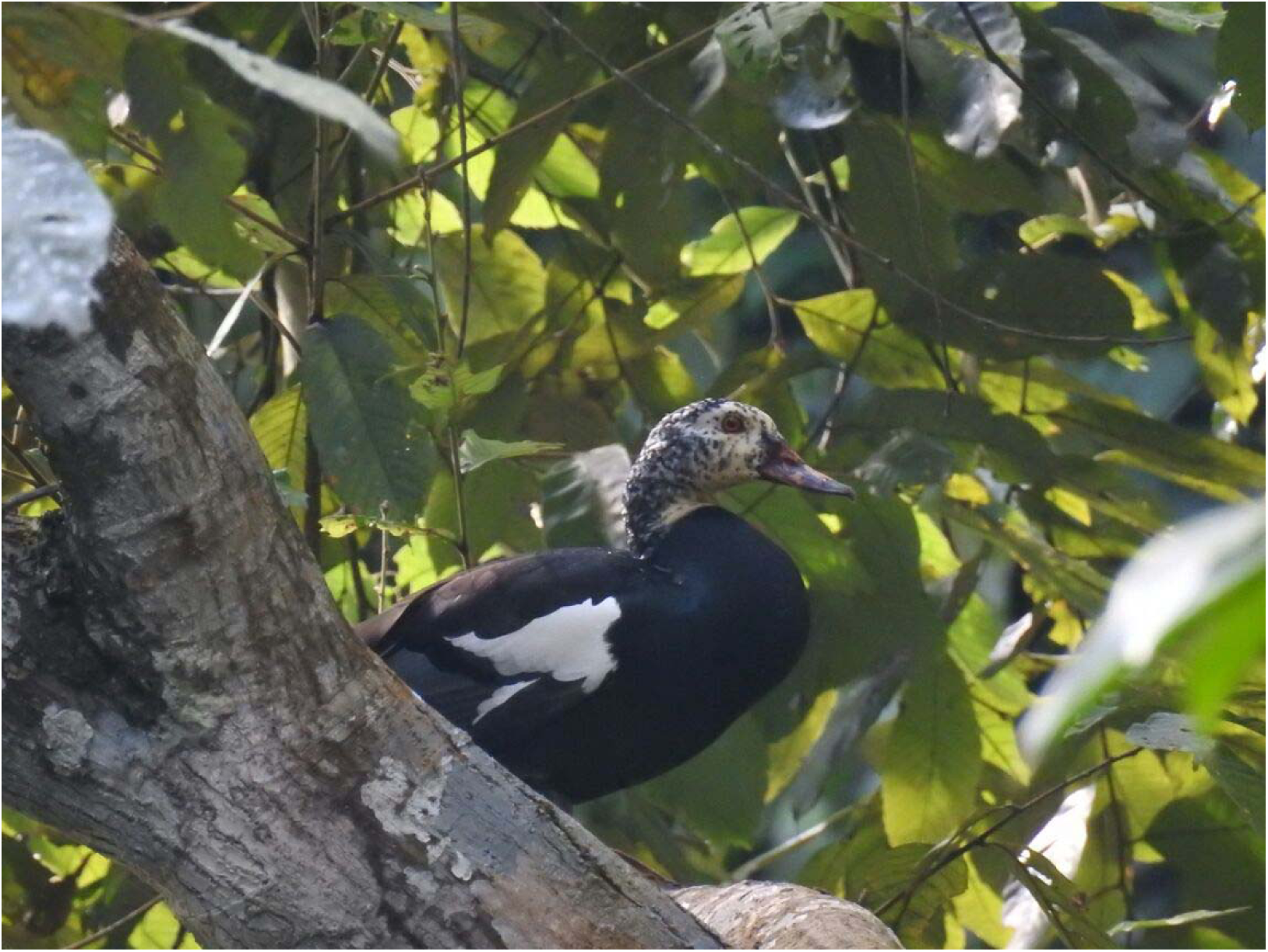
The white winged duck in its habitat in Dehing Patkai Wildlife Sanctuary (Credit: Binanda Hatibaruah)

Future optimal survey designs: Using standard design following MacKenzie and Royle (2005), we find that optimal number of repeated visits would have to be 4 and the optimal number of sites to be surveyed 524. On the other hand, using conditional occupancy design as proposed by Specht et al (2017), we estimated the number of optimal surveys to be 6 for future occupancy surveys, considering that the time and cost of different surveys remain the same.

## Discussion

Conservation threats to the white winged duck populations are manifold; they are threatened owing to rainforest habitat destruction, especially disturbances in riparian forests (Selvan et al. 2013). Selvan et al (2013) state that the survival of the species depends on the protection of dense and undisturbed primary rainforests. Sharma et al (2015) also point out that local populations of the duck are prone to local extinction. They are highly sensitive to habitat disturbance and human presence (Mackenzie and Kear, 1976) and hence chronic extraction of forest resources in the form of fuel wood and non-timber forest products (NTFPs) may have negative influences (Sharma et al. 2015). However, the degree to which the species is vulnerable to such threats is also unknown. The white winged duck is presently categorized as endangered (BirdLife International 2017), and listed under Schedule I of the Indian Wild Life (Protection) Act, 1972 (amended till date). It is also the state bird of Assam in India. However, the lack of basic ecology, population and threat status endangers the species greatly. Research on the species has been fairly neglected and the primary rainforests that they dwell in, in extension. Previous studies conducted have followed different methods and hence the change in abundance is yet to be confirmed. There is a need for researchers to agree upon and follow similar designs so as to provide better and real ground estimates of abundances and change in population trend, if any.

White winged duck occupancy was found to be low, however we acknowledge there were limitations to our survey. This study was conducted in only one season hence the interpretations of the models should be limited to just post monsoon season. Similar to earlier studies (Green 1993), we also found that the duck species is found in wetlands amidst forests. We identify such forest-wetland complexes as vital habitats for the white winged duck. These wetlands are usually surrounded by large trees. In Dehing Patkai, these large tree species were *Dillenia indica, Mesua ferrea, Shorea assamica, Vatica lancefolia* and *Dipterocapcus macrocarpus*. These wetlands were sometimes meshed with bamboo thickets as well. Our results also imply that low-lying wetlands with higher tree cover, along with the presence of dead logs (snags) in the wetlands are vital for white winged duck presence. Large trees contribute to high canopy cover which is important for this secretive bird. Fallen logs serve as perching stations for the duck.

This was our first attempt to understand the species occupancy in this rainforest area to aid conservation efforts. The study has generated fundamental understanding on the species occupancy and habitat ecology and explored on the design and methods needed to come up with better estimates of occupancy and detection. We hope that this will serve as a guiding report for future surveys for similar species. We recommend using occupancy models to better understand species ecology with the following important aspects to keep in mind while designing field surveys:

a. Number of sites (s) and number of replicates (K) to be kept at the maximum considering budget allocation and field limitation.
b. Effort should be maximised to achieve higher detection probability; we recommend focussing on highly probable sites for surveys to maximise detection probability and minimise error (SE or CI). From our understanding of this particular study area, we recommend using discrete sites such as ponds, and stream sides as survey sites (Mackenzie et al. 2003) rather than following a standard grid design. Higher effort in probable sites (targeted sampling; Harris et al. 2013) will provide robust estimates of *p* and hence *ψ*. Although grid based designs in wildlife surveys are used conventionally and often thought of as customary, Harris et al. (2013) found that targeted site sampling provide more precise estimates and indices of abundance. Harris et al (2013) further advocate the use of targeted sampling for species that have higher concentrations at resource concentrations. As per our occupancy models, we found higher occupancy in wetlands with high tree cover and lower elevation (low-lying ponds). Hence white winged ducks are concentrated and hence a targeted sampling is fit.
c. Although we explored two designs for future surveys, we suggest that using the conditional occupancy design (which would enable to survey more sites in limited time and survey attempts) would be optimal to understand space use intensity and species-habitat relationships. On the other hand, the standard design will be a better fit to study the abundance, true occupancy and population trends

We reiterate that a comprehensive and replicable field and analytical protocol is essential for better understanding the species’ ecology. Hence, we recommend focussed and long-term research and conservation efforts to protect and conserve the species.

## Author contributions

MP conceived the idea and acquired funding for the study. MP and SNJ deigned the study. MP collected all field data. SNJ analysed the data and led the manuscript preparation. LC and KC supervised the study and provided vital inputs during the field data collection. All authors approved the final draft of manuscript

## Acknowledgements

The study was part of a larger project entitled, ‘the white-winged duck: conservation status and site-specific threats to it in and around Dehing-Patkai Wildlife Sanctuary, Assam, India’. We thank the Assam State Forest Department for providing necessary permits to carry out field survey. We thank Dr Hannah Specht for providing useful comments/suggestions on various drafts of the manuscript. We also thank Genius, Dilip da and Bijoy da for their help during field work.

